# RSV & rhinovirus coinfection order dictates outcome via type III interferon-mediated interference in infant respiratory epithelium

**DOI:** 10.64898/2026.10.01.756024

**Authors:** Emily M. Schultz, Gina M. Aloisio, Ashley Murray, Sathi Wijeyesinghe, Letisha Aideyan, Trevor McBride, David Henke, Laura Ferlic-Stark, Gauri Khade, Oluwadamilola N. Oluwo, Hanning Li, Divya Nagaraj, Sarah E. Blutt, Vasanthi Avadhanula, Pedro A. Piedra

**Author notes:** **<u>Corresponding Author:</u>** Pedro A. Piedra, Baylor College of Medicine, 1 Baylor Plaza, Houston, TX 77030.

## Abstract

Coinfection with respiratory syncytial virus (RSV) and rhinovirus (RV) is the most frequently detected coinfection in infants, yet mechanisms governing viral interference and disease outcome remain poorly understood. Although infection order is a key determinant of coinfection outcome involving RSV, influenza, rhinovirus, and SARS-CoV-2, its role in RSV/RV coinfections has not been defined. Using infant-derived human nasal organoids at air-liquid interface, we demonstrate that coinfection order fundamentally determines the outcome of RSV/RV interactions. An initial RSV infection almost completely inhibits secondary RV coinfection, whereas primary RV infection only partially impairs a subsequent RSV coinfection. These distinct outcomes are accompanied by divergent temporal antiviral and pro-inflammatory responses. Mechanistically, we identify type III interferons as a sufficient mediator of RSV-induced restriction of RV, establishing a host-driven mechanism of viral interference. In contrast, partial impairment of RSV following RV infection occurs through an interferon-independent mechanism. Despite marked differences, RSV/RV coinfections do not exacerbate epithelial injury or innate immune responses beyond that induced by RSV alone, indicating RSV is the dominant driver of pathology in coinfections. Collectively, our data establishes coinfection order as a critical determinant of RSV/RV outcome and reveals novel and mechanistic insight into critical viral-host interactions in the infant nasal epithelium.

**Importance:** This study uncovers fundamental differences in infant epithelial control of two major pediatric viral pathogens. We demonstrate for the first time that the order of coinfection fundamentally determines the outcome of RSV-RV interactions within infant respiratory epithelium. By identifying type III interferon as an important regulator of these interactions, our study suggests that differences in interferon susceptibility among respiratory viruses can heavily influence coinfection outcomes. Importantly, despite clear differences in viral interference, RSV/RV coinfections did not produce greater epithelial injury or pro-inflammatory responses than RSV infection alone. Instead, RSV consistently emerged as the dominant driver of epithelial damage and inflammation, providing a mechanistic explanation for clinical observations that RSV/RV coinfections are often not more severe than RSV infection alone.

## Introduction

Respiratory viral infection is the leading cause of pediatric hospitalization in the US (1–3). Viral coinfections, most commonly between respiratory syncytial virus (RSV) and rhinovirus (RV), are detected in 30% of children (4–8). Although common, clinical reports remain divided if an RSV/RV coinfection increases ICU admission rates, oxygen supplementation, or length of hospitalization (9–12). Whether RSV/RV coinfection worsens clinical outcomes compared to single infection remains unresolved. One outpatient study in Chilean infants found no increase in severity with coinfection (9), while a multi-year prospective study in Spanish children reported elevated risk of fever and hypoxia (10). However, a key limitation of clinical studies is that they cannot determine the timing of viral infections-whether RV precedes RSV, RSV precedes RV, or both are acquired simultaneously.

Prior studies with other respiratory viruses have determined order dependent outcomes (13–18). In a human bronchial epithelial cell model, an initial RSV infection can protect against a severe sequential coinfection of severe acute respiratory syndrome coronavirus-2 (SARS-CoV-2) by activation of antiviral cytokines such as interferons (13). Alternatively, a primary infection of SARS-CoV-2 did not impair a secondary RSV coinfection. Similarly, interferon response from a primary RV infection has also been shown to impair a secondary coinfection of SARS-CoV-2, whereas a primary SARS-CoV-2 infection does not impact a secondary RV coinfection (16, 18). Whether order-dependent host innate immune responses and viral interference mechanisms impact RSV/RV coinfection dynamics remains unknown.

To address this gap, we utilized infant-derived human nasal organoids at air-liquid interface (HNO-ALI) as an ex vivo infant challenge model. HNO-ALIs are a critical and robust platform for studying RSV/RV coinfections since no animal models currently support simultaneous RSV and RV viral replication and disease recapitulation (19, 20). HNO-ALIs recapitulate the complex cellular composition of the upper respiratory epithelium and maintain the genetic background of their donor, allowing us to evaluate both viral and host responses. HNO-ALIs derived from infants compared to those derived from adults have increased susceptibility to RSV infection, increased cellular damage increased pro-inflammatory cytokines and type III interferons, establishing infant-derived HNO-ALIs as a valuable model for age specific co-infection question (21).

Here, we investigated RSV/RV coinfection in infant-derived HNO-ALIs to define how infection sequence influences virus-virus and host-virus interactions within the infant respiratory epithelium. We show that coinfection outcomes and the mechanisms governing viral interference are highly dependent on coinfection order. Primary RSV infection induces a robust type III interferon response that is sufficient to inhibit secondary RV coinfection, whereas RV-mediated restriction of RSV occurs independently of interferon signaling. We further identify RSV as the principal driver of epithelial injury and inflammatory responses during coinfection. These findings establish coinfection chronology as a key regulator of RSV-RV and host-virus interactions and provide insight into how early antiviral responses shape pathogen interactions in the infant respiratory epithelium.

## Results

### Alternate orders of RSV/RV coinfection differentially alter viral replication kinetics

To determine how coinfection sequence influenced viral replication, human nasal organoids at air liquid interface (HNO-ALIs) were infected with simultaneous coinfection, or alternate sequence of coinfection-RV preceding RSV (RV→RSV), or RSV preceding RV (RSV→RV) (FIG 1A). Single virus infections served as controls in addition to uninfected mock. In single virus infection, RV viral loads peaked 1-2 days post infection (dpi) and then declined to below the limit of detection (LOD) within 5-7 dpi (FIG 1B). In contrast RSV exhibited slower replication kinetics, with peak viral loads 3-4 dpi and maintained elevated levels through 10 dpi (FIG 1C).

**Figure 1.**
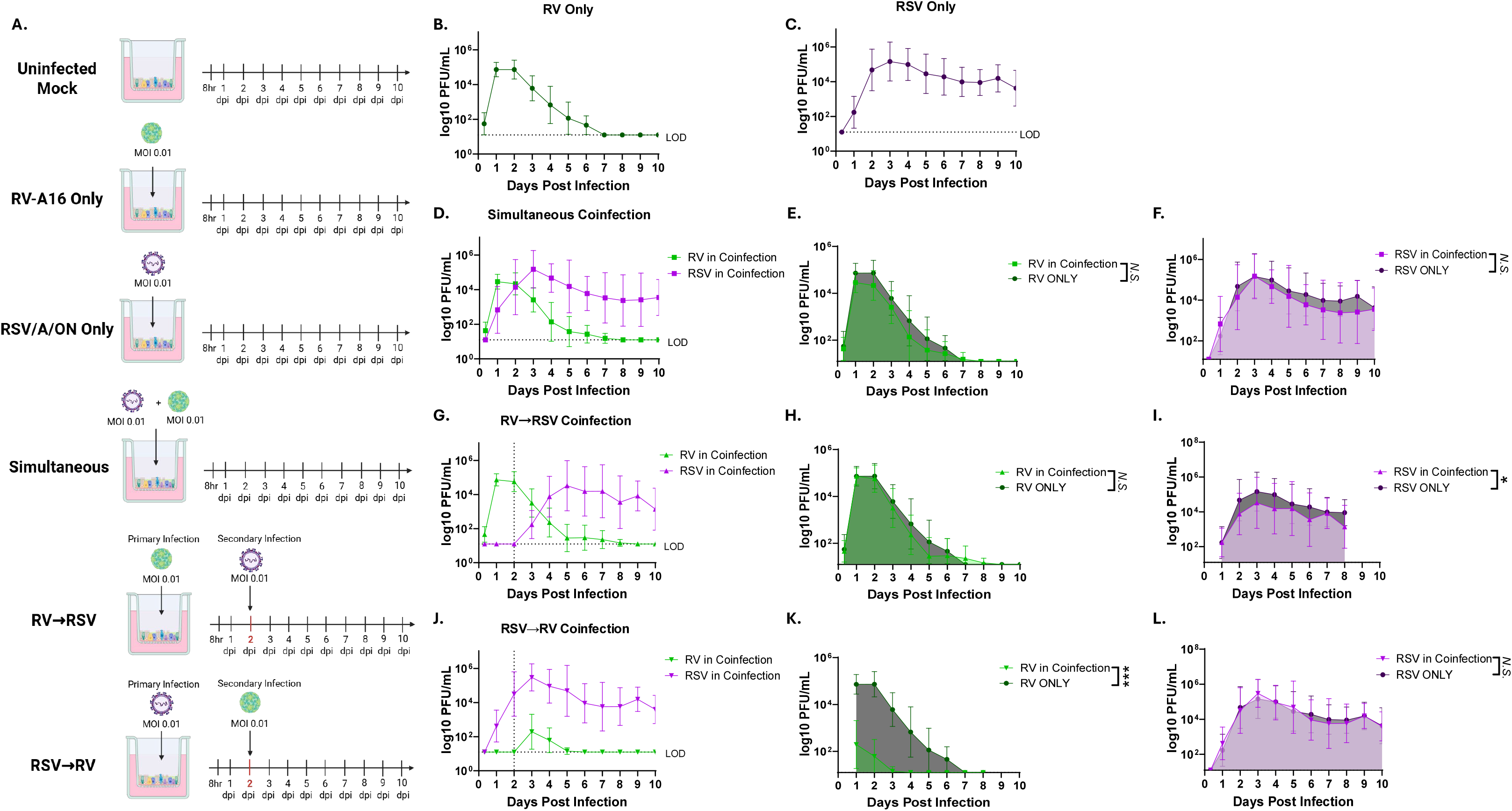
Different orders of coinfection alter viral replication kinetics. (A) Experimental design schematic. Viral quantification of **(B)** RV and **(C)** RSV single infections in infant HNO-ALIs. **(D)** Viral quantification of RV and RSV in simultaneous coinfection. **(E-F)** Area under the curve analysis of (E) RV viral load or (F) RSV viral load in a simultaneous coinfection compared to respective single virus control. **(G)** Viral quantification of RV and RSV in a RV→RSV sequence. **(H-I)** Area under the curve analysis of (H) RV viral load or (I) RSV viral load in a RV→RSV sequence compared to respective single virus control. **(J)** Viral quantification of RV and RSV in a RSV→RV sequence. **(K-L)** Area under the curve analysis of (K) RV viral load or (L) RSV viral load in a RSV→RV sequence compared to respective single virus control. All data displayed as average±SD of n=2 technical replicates from 5 different infant donor lines. Dotted y=1.096 indicates limit of detection (LOD), dotted x=2 line indicates day of coinfection. *Nonsignificant (N.S.), p>0.05*, \**p<0.05, **p<0.01, ***p<0.001* by Welch’s t-test.

In simultaneous coinfection, both RSV and RV replication kinetics were indistinguishable from their respective single-virus controls (FIG 1D). Consistent with this observation, area under the curve (AUC) analysis revealed no significant difference in cumulative viral replication for either RSV or RV relative to their respective single-virus infection (FIG 1E, F).

Sequential coinfection produced markedly different outcomes depending on which virus infected first. When RV preceded RSV (RV→RSV) (FIG 1G), RV replication was unaffected (FIG 1H). Whereas RSV viral load significantly dropped 56±35% when compared to RSV-only infection (FIG 1I). Despite this reduction, RSV replication remained detectable and relatively high (e.g. above 10^3^ on average) through 10 dpi.

In contrast, reversing the coinfection sequence (RSV→RV) resulted in near complete inhibition of RV (FIG 1J). AUC analysis revealed a significant 99.3±0.8% reduction in infectious RV compared to single RV control (FIG 1K). In contrast, RSV was not impacted and continued to replicate at sustained viral loads for 10 days (FIG 1L). These data demonstrate that a primary RSV infection in infant HNO-ALIs can severely inhibit a secondary RV coinfection when introduced 2 days later.

Together, our findings reveal that RSV/RV coinfection outcomes are highly dependent on infection sequence. While simultaneous coinfection had minimal impact on viral replication, sequential infection resulted in disproportionate viral interference. RSV exerted a substantially stronger inhibitory effect on a secondary RV coinfection compared to the impact RV had on a secondary RSV coinfection.

### RSV and RV coinfection does not result in increased cellular damage or cell death

Having established that infection order differentially affects viral replication kinetics, we next examined whether these virologic differences translated to corresponding changes in epithelial injury and cell death. Infant HNO-ALIs were processed for hematoxylin and eosin (H&E) staining at 2, 4, and 10 dpi. At 2 dpi all infection groups resembled mock infected cultures and retained an intact nasal epithelium with noticeable apical ciliated cells (FIG 2A). Histopathological changes remained minimal following RV infection, and by 10 dpi RV infected HNO-ALI closely resembled mock controls, consistent with the limited epithelial injury reported in RV infected adult biopsies (22) (FIG 2A). In contrast, RSV only, simultaneous, and RSV→RV coinfected HNO-ALIs at 4 dpi displayed rounded apical ciliated cells, a hallmark of RSV-induced epithelial injury and cell extrusion (21, 23) (FIG 2A). RV→RSV coinfected HNO-ALIs did not display this characteristic at 4 dpi but showed considerable damage at 10 dpi once RSV infection was established. By 10 dpi, RSV only and all coinfected conditions displayed widespread damage, characterized by loss of ciliated cells and epithelial sloughing (FIG 2A).

**Figure 2.**
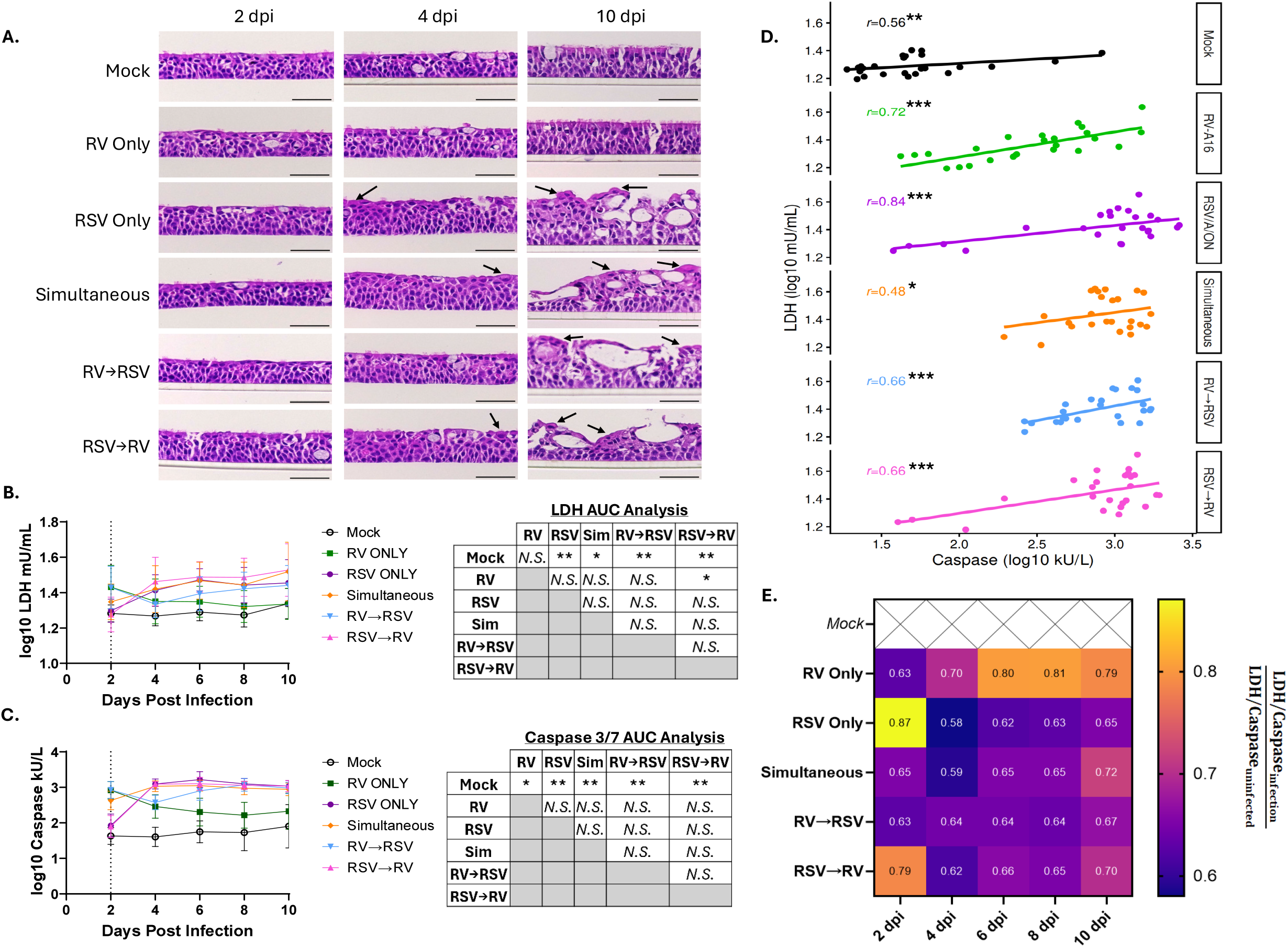
Different orders of coinfection do not result in increased cellular damage or apoptosis. **(A)** Representative images of hematoxylin and eosin stained donor-matched infant HNO-ALIs at 2, 4, and 10 days post infection (dpi) with either i) mock infected ii) RV only iii) RSV only iv) simultaneous RSV+RV coinfection v) RV→RSV coinfection or vi) RSV→RV coinfection. Scale bar= 50µm in all images and is located on the basolateral side of the HNO-ALI. Arrow indicates rounded cell pathology typical of RSV infection. **(B)** LDH concentration over time with AUC analysis **(C)** Caspase 3/7 concentration over time with AUC analysis. (B&C) Data displayed as average±SD, n=2 technical replicates from five different infant donor lines. Dotted x=2 line indicates day of coinfection for RV→RSV and RSV→RV groups. *Nonsignificant (N.S.), p>0.05*, \**p<0.05, **p<0.01, ***p<0.001* by Welch’s t-test. **(D)** LDH:Caspase repeated measures correlation *Nonsignificant (N.S.), p>0.05*, \**p<0.05, **p<0.01, ***p<0.001*. Data displayed as averaged n=2 technical replicates from five individual donor lines at five time-points. **(E)** Fold change LDH:Caspase ratio over time.

To quantitatively assess cellular injury and apoptosis, we measured release of lactate dehydrogenase (LDH) and caspase 3/7 activity within HNO-ALI samples. LDH release, a marker of membrane damage, and caspase 3/7 activity, a marker of programmed apoptosis, have both been linked to epithelial injury during viral respiratory infections and been used as biomarkers in clinical samples (24–28). RV infection induced early increases in both LDH and caspase 3/7 that peaked 2 dpi and subsequently declined toward baseline levels observed in mock HNO-ALIs (FIG 2B-C). Consistent with the limited histologic changes observed following RV infection, cumulative LDH release (AUC 5±5% increase relative to mock) was not significantly different from mock infected controls (FIG 2B). However, there was a modest 43±22% increase in caspase 3/7 activity in RV infected HNO-ALI compared to mock infected (FIG 2C). In contrast, RSV infection induced a sustained 11±5% increase in LDH and 80±34% increase in caspase activity compared to mock infected controls (FIG 2B-C). LDH and caspase 3/7 levels parallelled viral replication kinetics, with maximal responses occurring 4-10 dpi (FIG 1C, 2B-C). Similarly, all coinfection conditions exhibited significantly greater cumulative LDH release (Simultaneous: 11±7%, RV→RSV: 8±4%, RSV→RV: 13±5%) and caspase 3/7 activity (Simultaneous: 79±33%, RV→RSV: 74±32%, RSV→RV: 77±32%) compared to mock controls (FIG 2B-C). However, neither simultaneous coinfection nor sequential coinfections resulted in greater cumulative LDH release or caspase 3/7 activity than RSV infection alone. There were also no differences in cumulative cellular injury or apoptosis among the coinfection groups (FIG 2B-C). Furthermore, LDH and caspase 3/7 concentrations were significantly and positively correlated across all infection conditions (r = 0.48–0.84) (FIG 2D). This indicates that increased epithelial injury was generally accompanied by increased apoptosis, and different coinfection sequences did not disrupt this association.

To quantify the relationship between LDH and caspase during coinfection, we calculated the fold change in LDH:caspase ratio relative to mock infected controls. LDH:caspase ratios in clinical samples can be predictive of outcome in children, with a higher ratio being associated with hospitalization (27). All infection conditions exhibited reductions in the LDH:caspase ratio, indicating a shift toward greater caspase activity relative to epithelial injury (FIG 2E). RV infection induced an early reduction in the LDH:caspase ratio that progressively returned toward baseline 6-10 dpi, consistent with viral clearance. In contrast, RSV infection did not induce a marked reduction in the LDH:caspase ratio until 4 dpi with a ratio of 0.58 and remaining between 0.62-0.65 through 10 dpi (FIG 2E). Simultaneous and sequential coinfection exhibited similar sustained reductions, closely resembling an RSV only infection. Notably, the LDH:caspase ratio remained lower despite increasing LDH release in RSV-co**n**taining conditions, demonstrating that increases in apoptosis outpace increases in cellular injury (FIG 2B, C, E). Together, these data indicate that although epithelial injury and apoptosis remain positively associated, RSV containing infections exhibit a disproportionately greater increase in apoptosis relative to cellular injury.

Collectively, histopathology, LDH release, caspase 3/7 activity, and LDH:caspase dynamics revealed a consistent pattern across experimental groups. RSV-containing infections produced the most sustained epithelial damage and apoptosis, while RV infection induced a comparatively transient response. These findings are consistent with histopathological and clinical reports demonstrating that RSV induces substantially greater epithelial injury than RV (29–31). Despite the distinct viral replication kinetics observed among coinfection conditions, neither simultaneous nor sequential coinfections increased epithelial injury or apoptosis beyond levels observed during RSV infection alone. These findings indicate that RSV is the primary driver of epithelial cell death dynamics during RSV/RV coinfections in infant HNO-ALI.

### Epithelial cytokine and chemokine response is driven by RSV in different orders of coinfection

To evaluate the infant epithelial innate immune response during different coinfection sequences, we quantified 12 distinct cytokine and chemokines at 2, 4, and 6 dpi in infant HNO-ALIs. Cytokine and chemokine expression were normalized as z-scores and analyzed individually and by functional group relative to mock infected controls using adjusted odds ratios.

Overall cytokine and chemokine production was similar across all infection conditions at 2 dpi (FIG 3A, Supp Data). However, innate immune responses diverged substantially at 4 dpi. Groups containing four days of RSV replication (RSV only, simultaneous, or RSV→RV) exhibited greater overall cytokine production compared to RV only infection (FIG 3A). This pattern persisted through 6 dpi when all RSV-containing groups displayed significantly increased cytokine and chemokine production relative to RV infection alone (FIG 3A, Supp Data). Despite concurrent replication of both viruses, simultaneous coinfection did not enhance cytokine expression beyond levels observed during RSV only infection. This indicates that sustained epithelial immune activation was not driven by the presence of two replicating viruses but instead reflects the dominant influence of RSV replication in infant respiratory epithelium.

**Figure 3.**
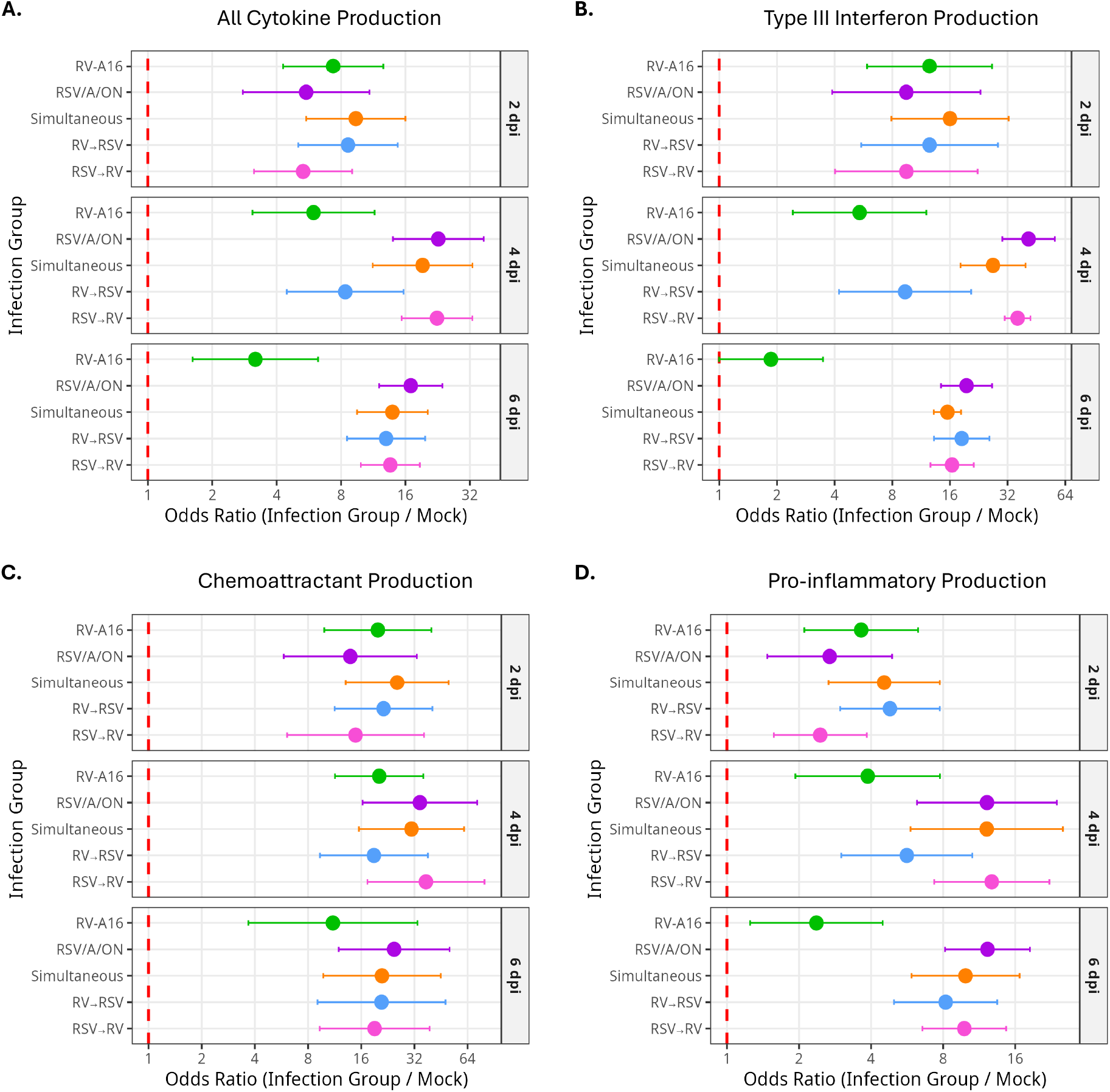
Epithelial cytokine and chemokine release is driven by RSV in different orders of coinfection. Forest plots of the adjusted odds ratio compared to mock with 95% confidence interval of all infection groups at 2, 4, and 6 days post infection (dpi) for functional groups: **(A)** Overall cytokine production **(B)** Type III interferon production (comprised of IFN-λ1, IFN-λ2, and IFN-λ3) **(C)** Chemoattractant production (comprised of Eotaxin and IP-10) **(D)** Pro-inflammatory cytokine production (comprised of TNFα, IL-1α, IL-1ß, IL-6, and IL-8). Data analyzed from n=2 technical replicates and five infant donor lines biological replicates. Significance indicated by non-overlapping confidence intervals. See also Figure S1, S2, Supp Data.

Type III interferons (IFN-λ1, IFN-λ2, IFN-λ3) represented the predominant antiviral response in infected infant HNO-ALIs. In contrast, type I interferons (IFN-α2 and IFN-ß) remained undetectable throughout infection. By 4 dpi, RSV only, simultaneous, and RSV→RV infection groups produced significantly greater amounts of type III IFNs compared to RV infection alone (FIG 3B). Elevated production persisted through 6 dpi, when all RSV containing groups exhibited a significantly greater type III IFN response than RV only infection (Supp Data). Notably, the reduction in RSV viral load observed in RV→RSV coinfection (FIG 1F) was accompanied by a corresponding decrease in peak IFN-λ production (Supp Data), linking epithelial type III interferon responses to RSV replication. Consistent with clinical observations of RSV- and RV-associated bronchiolitis (32), peak IFN-λ production during RV infection remained substantially lower than that induced by RSV infection (FIG 3B, Supp Data). Together, these findings identify RSV replication as the primary determinant of type III interferon production during RSV/RV coinfection.

Although individual IFN-λ subtypes (IFN-λ1, IFN-λ2, IFN-λ3) exhibited similar kinetics across infection groups (FIG S1A-C), substantial donor-to-donor variation was evident. Individual HNO-ALI donor lines consistently favored expression of a specific IFN-λ subtype irrespective of the infecting virus (FIG S2), suggesting intrinsic epithelial programs influence the composition of type III IFN responses. These observations underscore the importance of evaluating multiple donor lines and measuring all IFN-λ subtypes when characterizing epithelial antiviral immunity.

We next examined epithelial chemoattractant responses via chemokines IP-10 and Eotaxin. IP-10 is a potent chemoattractant for T cells and NK cells and has been implicated with increased severity during respiratory virus infection(33, 34). Eotaxin attracts eosinophils and has been associated with RV induced airway inflammation and allergic airway disease (35–37). Despite their distinct immunological functions, neither the overall chemoattractant response nor individual production differed significantly among RSV, RV, or coinfection conditions at any time point examined (FIG 3C & FIG S1D-E). In contrast to other analytes tested, RV only, RSV only, and all coinfection conditions produced similar elevated levels of IP-10 (FIG S1E). Thus, unlike type III IFN responses, epithelial chemokine induction appeared largely independent of viral identity and was not augmented by alternate coinfection sequences.

We also evaluated pro-inflammatory cytokines, including tumor necrosis factor alpha (TNFα), interleukin 1 alpha (IL-1α), interleukin 1 beta (IL-1ß), interleukin 6 (IL-6), and interleukin 8 (IL-8). Elevated pro-inflammatory cytokines have been found in pediatric nasal washes during severe RSV and/or RV infection (37, 38). Similar to type III IFNs, pro-inflammatory responses closely paralleled RSV replication kinetics. At 4 dpi, RSV-only, simultaneous coinfection, and RSV→RV cultures exhibited elevated pro-inflammatory cytokine production, whereas RV-only and RV→RSV cultures remained near baseline levels (FIG 3D). By 6 dpi, all RSV-containing groups displayed significantly greater pro-inflammatory cytokine production than RV only infection (FIG 3D, Supp Data). Peak pro-inflammatory responses during RV infection remained substantially lower than those observed during RSV infection, further supporting a dominant role for RSV in shaping epithelial innate immune responses (FIG 3D, Supp Data).

Collectively, our data demonstrates that RSV elicited a stronger and more sustained epithelial innate immune response over 10 days compared to RV in infant respiratory epithelium at the same moi. Across all coinfection conditions, the magnitude and kinetics of both type III interferon and pro-inflammatory cytokine production closely track RSV replication dynamics. Importantly, simultaneous coinfection did not amplify innate immune responses beyond levels observed during RSV only infection. Together, these findings identify RSV replication as the principal driver of epithelial immune responses during RSV/RV coinfection, regardless of coinfection order.

### Type III interferons selectively inhibit rhinovirus but not RSV replication

Given the greater induction of type III IFNs during RSV infection, we postulated that type III IFNs contribute to the viral interference observed during sequential RSV/RV coinfections. To model the anti-viral state established by a prior viral infection, recombinant human interferon (rhIFN-λ1, rhIFN-λ2, or rhIFN-λ3) was added to the basolateral compartment of infant HNO-ALIs 24 hours prior to viral infection. Biological activity of each rhIFN-λ was confirmed by induction of interferon stimulated gene *MX1* (FIG S3C-E).

Pretreatment with individual rhIFN-λ1, rhIFN-λ2, or rhIFN-λ3 at 100ng/mL, a concentration comparable to peak levels previously detected in RSV infected HNO-ALIs, potently inhibited RV but not RSV. Relative to untreated controls, AUC analysis revealed RV replication was reduced by 93±5%, 90±11%, and 95±3% following treatment with rhIFN-λ1, rhIFN-λ2, rhIFN-λ3 respectively (FIG 4A). In stark contrast, pretreatment with any individual IFN-λ subtype did not have any detectable effect on RSV viral loads (FIG 4B). A 10-fold dose dependent response analysis demonstrated similar antiviral potency among all three IFN-λ subtypes, with each producing similar concentration-dependent inhibition of RV replication (FIG S4 A-C).

**Figure 4.**
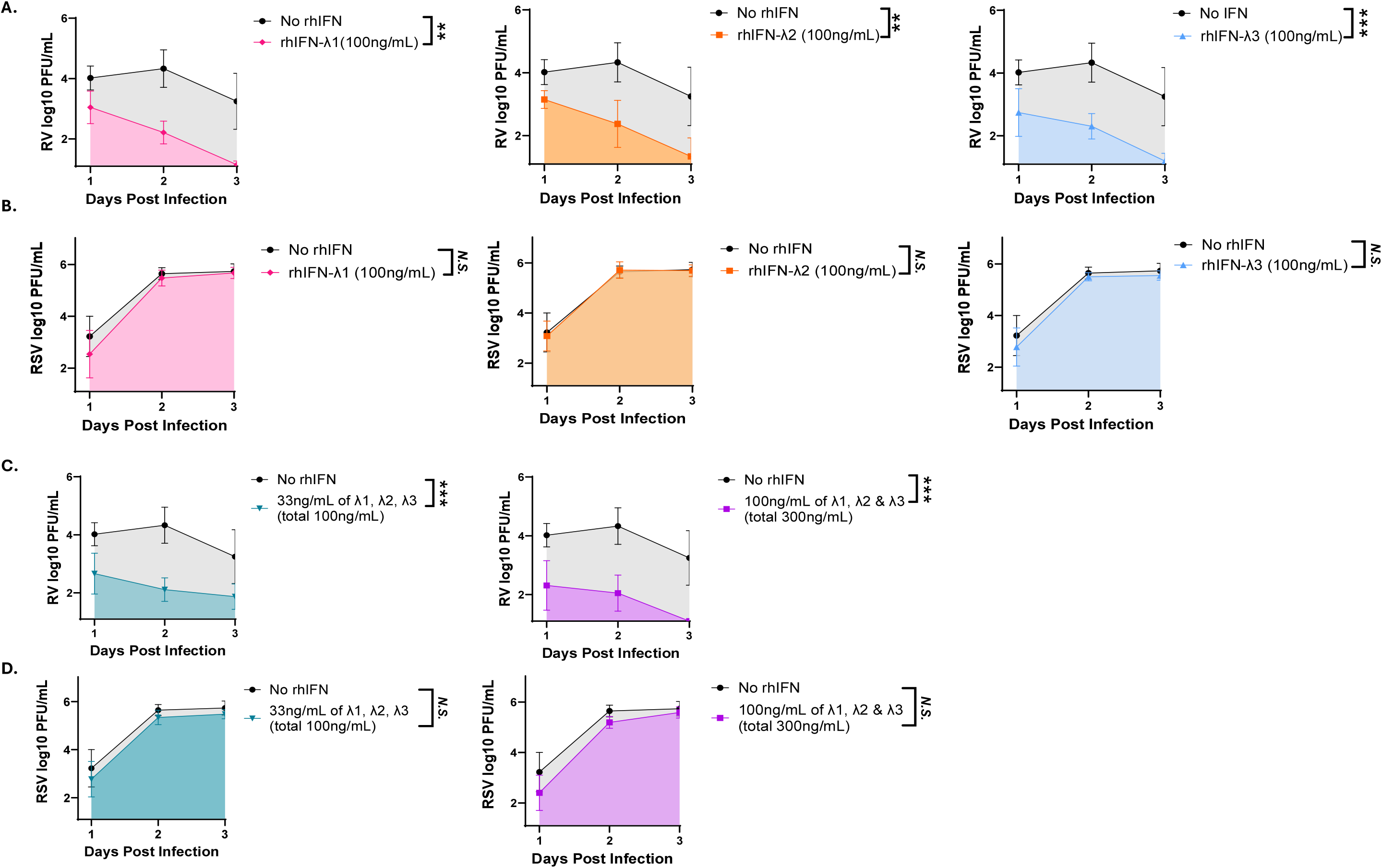
Type III interferons inhibit rhinovirus replication but not RSV replication. **(A)** RV or **(B)** RSV replication over time in the presence of 100ng/mL of either rhIFN-λ1, -λ2, or -λ3 along with corresponding area under the curve analysis. **(C)** RV or **(D)** RSV replication over time in the presence of either 33ng/mL of each rhIFN-λ1, -λ2, and -λ3 (total 100ng/mL) or 100ng/mL of each rhIFN-λ1, -λ2, or -λ3 (total 300ng/mL) along with corresponding area under the curve analysis. All data displayed as average±SD of n=2 technical replicates from 3 different infant donor lines. *Nonsignificant (N.S.) p>0.05*, \**p<0.05, **p<0.01, ***p<0.001* by Welch’s t-test. Also see Figure S3, S4.

Because all three type III IFNs are produced by the respiratory epithelium during viral infection, we next assessed whether combined exposure to rhIFN-λ1, rhIFN-λ2, and rhIFN-λ3 enhanced antiviral activity. HNO-ALIs were incubated with either low-dose (33.3 ng/mL per subtype; 100 ng/mL total) or high-dose (100 ng/mL per subtype; 300 ng/mL total) IFN-λ mixtures 24 hours prior to infection. Both low and high dose combinations robustly suppressed RV replication by 97±2% and 96±3% respectively (FIG 4C). However, neither combination produced greater inhibition than individual IFN-λ subtype treatment (FIG S3A), indicating that IFN-λ subtypes do not exhibit additive or synergistic antiviral activity in respiratory epithelium at the doses tested. Moreover, consistent with the individual IFN-λ subtypes treatments, neither IFN-λ combination significantly altered RSV replication (FIG 4D, S3B).

We propose that type III IFNs are a sufficient mechanism for how RSV can severely impair a secondary coinfection of RV in an RSV→RV order of coinfection (FIG 2F). In contrast, since RSV is not impaired by the presence of type III IFNs, this is not a mechanism used by RV to impair a secondary coinfection of RSV in a RV→RSV order of coinfection (FIG 2C).

Together, these findings demonstrate that type III interferons selectively restrict RV, not RSV, replication in infant airway epithelium. This selectivity offers a mechanistic basis for the near-complete suppression of secondary RV infection in the RSV→RV coinfection setting, where RSV-induced IFN-λ precedes RV exposure. Conversely, because RSV replication was resistant to IFN-λ-mediated restriction, type III interferons alone are unlikely to explain the reduced RSV replication observed during RV→RSV coinfection.

### Blocking interferon lambda receptor 1 allows for increased rhinovirus replication but has no impact on RSV replication

To further validate that type III IFN signaling is required for the viral interference observed during RSV→RV coinfection, we blocked the shared receptor subunit for all IFN-λ subtypes, interferon lambda receptor 1 (IFNLR1). Infant HNO-ALIs were treated with a monoclonal antibody against IFNLR1 or an IgG isotype control 24 hours prior to infection with either RV or RSV (FIG 5A). Blockade of IFNLR1 significantly increased RV replication by 638±104% relative to isotype-treated control (FIG 5B). This demonstrated that endogenous type III IFN signaling restricts RV replication in infant respiratory epithelium. RV replication, however, was not completely unimpaired when blocking IFNLR1, with viral titers decreasing below peak values after 3 dpi (FIG 5B). This observed decrease in RV replication, while remaining elevated compared to the IgG control, is likely due to noncanonical induction of interferon stimulated genes during viral infection, evidenced by *MX1* induction (FIG S5) (39). In contrast, IFNLR1 blockade had no effect on RSV replication (FIG 5C), further supporting the conclusion that RSV is resistant to IFN-λ mediated interference.

**Figure 5.**
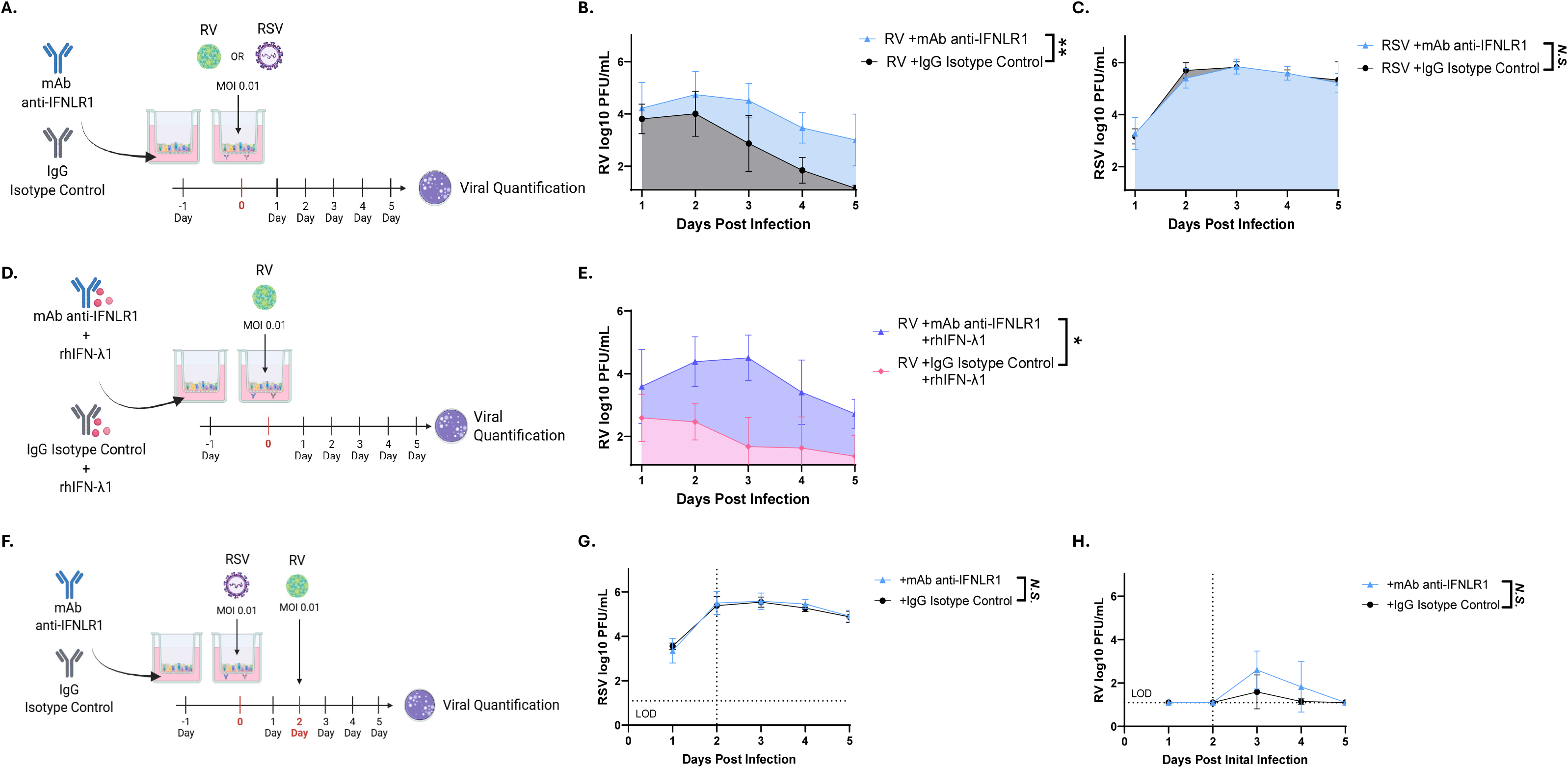
Blocking interferon lambda receptor 1 allows for increased rhinovirus replication but has no *impact on RSV replication.* **(A)** Experimental design schematic for IFNLR1 blocking experiment during RV or RSV infection. **(B-C)** AUC analysis of (B) RV or (C) RSV replication over time in presence of anti-IFNLR1 mAb. **(D)** Experimental design schematic for RV replication when blocking signaling of high exogenous rhIFN-λ1 through IFNLR1. **(E)** AUC analysis of RV replication over time when blocking exogenous levels of rhIFN-λ through IFNLR1. **(F)** Experimental design schematic of RSV→RV coinfection when blocking the IFNLR1 receptor. **(G)** AUC analysis of RSV replication over time when blocking signaling through IFNLR1 during a RSV→RV coinfection. **(H)** AUC analysis of RV replication over time when blocking signaling through IFNLR1 during a RSV→RV coinfection. All data displayed as average±SD of n=2 technical replicates from 3 different infant donor lines. Dotted y-line indicates limit of detection (LOD), dotted x=2 line indicates day of coinfection. *Nonsignificant (N.S.) p>0.05*, \**p<0.05, **p<0.01, ***p<0.001* by Welch’s t-test. Also see Figure S5.

Because RSV induced substantially greater IFN-λ production than RV infection alone (FIG 3B, S1A-C), we next tested whether IFNLR1 blockade could restore RV replication in the presence of high exogenous IFN-λ (FIG 5D). Anti-IFNLR1 treatment effectively inhibited IFN-λ signaling (FIG S5) and significantly rescued RV replication compared to cultures not treated with anti-IFNLR1 (FIG 5E). These findings further support type III IFNs as a sufficient mechanism for suppressing RV replication.

To determine the impact of IFNLR1 during RSV→RV coinfection, we once again pre-treated infant HNO-ALIs with either anti-IFNLR1 mAb or IgG isotype 24 hours prior to coinfection (FIG 5F). Consistent with previous experiments, IFNLR1 blockade did not alter RSV replication (FIG 5G). Although RV replication increased following IFNLR1 blockade (average 1239-fold increase), the magnitude of rescue varied considerably among donor lines and did not reach statistical significance (FIG 5H). Expression of interferon stimulated gene *MX1* remained elevated during viral infection despite IFNLR1 blockade (FIG S5). *MX1* levels were similar to isotype treated controls, indicating noncanonical induction of interferon stimulated genes in the absence of interferon lambda signaling (39).

Overall, these data demonstrate that type III interferons are sufficient to inhibit RV replication but are not strictly necessary for inhibition of RV during RSV→RV coinfections. In contrast RSV replication remained unaffected by IFNLR1 blockade. This indicated that RSV is intrinsically resistant to IFN-λ-mediated interference and that an alternative mechanism likely contributes to reduced RSV replication during RV→RSV coinfection. All together this data shows that type III IFNs are a sufficient mechanism to impair RV replication but not necessary for interference during an RSV→RV coinfection.

## Discussion

Using infant-derived HNO-ALI cultures, we show for the first time that the outcomes and mechanisms governing viral interference in RSV/RV coinfection are highly dependent on coinfection order. We found that a primary infection of RSV can cause near complete inhibition of a secondary RV coinfection, and we demonstrate that type III interferons (IFNs), provoked by a primary RSV infection, as a major mechanism of RV interference. We also observed significant impairment of RSV replication when it followed a primary infection of RV. Exogenous type III IFNs had no impact on RSV replication, suggesting an interferon-independent interference mechanism that remains to be defined. Our ex vivo infant study also mirrors multiple clinical studies showing that RSV/RV coinfections do not result in more severe damage and pro-inflammatory response compared to RSV infection alone (31, 38, 40). We propose that no matter the coinfection order, RSV is the main driver of nasal epithelial responses in RSV/RV coinfection.

The replication kinetics of RSV and RV show distinct trajectories in our ex vivo infant challenge HNO-ALI model. RV is inherently controlled and cleared from the nasal epithelium anywhere from 5-8 dpi. In contrast RSV replication remains at elevated infectious levels through 10 dpi. Our observed interference between RSV and RV also agrees with a study from Essaidi-Laziosi et al. in primary airway epithelial cells (41). Essaidi-Laziosi et al. however, did not observe impairment of RSV after a primary infection of RV (RV→RSV coinfection order). This non-concordance in results could be confounded by differences in study design. Essaidi-Laziosi et al. only used one donor line of unknown donor age an is unable to account for potential donor variability, which was observed within our study (41). Further the study quantified genomic material at one-time point post coinfection, potentially missing the temporal relationships of viral load across time in a coinfection (41). To our knowledge, this immense control and clearance of RV replication has not been shown before. Many previous studies on RV replication were performed in animals, homogenous immortalized cell culture, or infection in primary epithelial cell cultures that were not studied beyond 4-5 dpi (42–45). Additionally, the persistence of RSV in infant HNO-ALIs closely mimics the prolonged shedding seen in immunocompromised children unable to resolve infection, especially those with CD4+/CD8+ T cell deficiencies (46–48). Our findings highlight that innate immune factors orchestrated from epithelium alone are sufficient to control RV infection but not for RSV where a secondary cellular immune response is likely needed to control and clear infection.

Simultaneous coinfection of RSV and RV had no impact on the replication of either virus. However, in sequential coinfections (either RV→RSV or RSV→RV) the second infecting virus was significantly impaired, with greater impact on viral load reduction for RV (99%) in an RSV→RV coinfection compared to RSV (56%) in an RV→RSV coinfection. The observed near complete inhibition of RV after a primary infection of RSV is consistent with clinical birth cohort studies that show decreased odds of having a RV infection after testing positive for RSV (49), suggesting that RSV infection can prevent RV infection. Epidemiological surveillance data showing a decrease in RV cases during periods of high RSV positivity within communities (50) further supports the concept of viral interference, however at the population level. Taken together this suggests that most common RSV/RV coinfections in infants are most likely either a simultaneous or RV→RSV coinfection order where both viruses can co-exist, and not an RSV→RV coinfection order where RV is almost completely blocked.

We also found evidence that cytopathology within RSV/RV coinfections is driven by RSV, not RV, and not impacted by coinfection order. H&E-stained cross-sections of HNO-ALIs coinfected with alternate orders all resembled an RSV only infection by 10dpi with increased epithelial damage. Furthermore, markers of cellular injury (LDH) and apoptosis (caspase 3/7) indicated no significant differences between different orders of coinfection and an RSV only infection. This matches previously published clinical findings that LDH and caspase concentrations in nasal wash samples are the same from children in the emergency room with either RSV/RV coinfection or single RSV or RV infections (27). Furthermore, clinical reports also show that RSV-bronchiolitis is more severe than RV-bronchiolitis in hospitalized infants and that RSV can be the driver of severity in RSV/RV coinfections (4, 8, 31, 40). Consistent with this, RSV only and RSV/RV coinfected infant HNO-ALIs exhibited a more pronounced and sustained pro-inflammatory cytokine profile through 10dpi compared to RV infected cultures. This pattern in RSV/RV coinfected conditions could be due to the lack of infectious RV by 10 dpi. These findings are consistent with clinical studies showing that elevated levels of the pro-inflammatory cytokines IL-6, IL-1ß, and IL-8 findings in children with RSV only or RSV/RV coinfection are associated with increased disease severity and greater need for supplemental oxygen (38). Collectively, these data suggest that RSV is driver of severity and inflammation, and we can therefore speculate that severity observed in RSV/RV coinfections is primarily driven by RSV, rather than an additive effect contributed by RV.

Our study found a unique pattern of viral inhibition with type III IFN unable to impair RSV replication yet inhibiting RV replication 90-95% within infant HNO-ALIs. Interferons have been known to impair respiratory viruses like RV and RSV since the 1980s (51–53). However, many of these early studies in either a human challenge model or immortalized cell culture solely utilized type I IFNs since type III IFNs were not described until 2003 (54). Type I and type III IFNs induce similar sets of interferon stimulated genes (ISGs), but there is a temporal difference in expression, creating distinct antiviral response profiles (55). The ability of type III IFN produced ISGs to have a specific viral interference impact on RV and not RSV could point towards a potential targeted biotherapeutic for RV infection in infants. Although RV replication was not completely rescued in RSV→RV coinfection when blocking IFNLR1, the specific ISGs causing interference were likely still produced via noncanonical and redundant pathways for when IFN signaling fails (39). Delineating what specific ISG present inhibits RV-A16 replication in infant respiratory epithelium could prove a valuable addition to the field since there are over 160 different types of RV, complicating the development of a broad acting vaccine (56, 57).

Given that type III IFN had no impact on RSV replication in the infant respiratory epithelium in our infant HNO-ALI model, further investigation is needed to interrogate the observed impairment on RSV replication in an RV→RSV coinfection. Both RSV and RV primarily target and replicate within apical ciliated cells of the respiratory epithelium (58, 59). This could create competition for susceptible cell populations during a coinfection. Increased caspase 3/7 activity and increased TNFα and IL-1ß cytokines connected to apoptosis after RV infection potentially point towards a reduced susceptible cell populations when RSV is introduced 2 days post RV infection. This impaired availability of susceptible cells could serve as a mechanism for how RV interferes with a secondary coinfection of RSV (RV→RSV). Alternatively, other pathways associated with viral replication that affect interferon-independent ISG response remain poorly understood. These potential viral interference pathways impacting RSV should be explored further.

While our study provides a detailed and mechanistic account of alternate orders of RSV/RV coinfection within the infant respiratory epithelium, it is limited in its interpretation. HNO-ALI cultures, while recapitulating the cellular composition of the upper respiratory tract, is an epithelial only model system without any cellular innate or adaptive immune responses. The lack of tissue-resident memory T cells, neutrophils, NK cells, monocytes, and other immune cell populations that could have an impact on viral control, cellular damage, and the cytokine profiles observed. Further iterations of HNO-ALI culture systems containing immune cell populations are needed to better understand their impact on the respiratory epithelium and coinfection states. Furthermore, the respiratory epithelium is a dynamic site with microbial communities that have the potential to influence viral infection (60, 61). The HNO-ALI system used in this study did not include any bacterial colonization.

An additional limitation of this study is that only one strain of RSV (RSV/A/ON) and RV (RV-A16) were utilized. RSV/A/ON was selected because of being the main contemporaneous RSV-A strain circulating. Strains from both RSV A and B subgroups can cause severe disease. There has not been substantial evidence to point towards clinical differences between the two. RV-A16 was selected due to being of the major receptor class for rhinoviruses-intercellular adhesion molecule 1 (ICAM-1) (62). While there is immense genetic variation in rhinoviruses, most circulating strains are either RV-A or RV-C with both being linked to more severe respiratory infections (56). Additionally, recent sequencing analysis for the past 3 decades has shown RV-A16 remains relevant circulating strain (57). Even though only one type of RV and RSV were used in this analysis, we believe the results to be broadly applicable to other RSV and RV types due to strategic selection of representative types. Furthermore, with the establishment of a model system, further factors such as additional viral subtypes, different time points of coinfection, and different multiplicities of infection can be evaluated.

In summary, our study harnessed infant HNO-ALIs to find that the order of RSV/RV coinfection dictates temporal host: virus interactions and outcomes. We display that a primary infection of RSV can cause near complete inhibition of a secondary RV coinfection, suggesting that RSV/RV coinfections detected in hospitalized infants are not of the RSV→RV order, and likely a RV→RSV or simultaneous coinfection order. Further mechanistic investigation found type III IFN to be a unique viral interference mechanism for how RSV inhibits a secondary coinfection of RV, yet not the interference mechanism involved in the alternate coinfection order of RV→RSV. Furthermore, RSV was observed to be the main driver of epithelial damage and the production of pro-inflammatory cytokines connected to increased cytopathogenicity. Coinfected infant HNO-ALIs recapitulate viral pathology and host responses as previously reported in clinical samples from hospitalized infants, establishing a reliable model system for investigating RSV/RV coinfections and viral interference mechanisms. We believe this study provides foundational knowledge to build upon for better understanding the most common coinfection found in infants hospitalized for severe respiratory distress. Understanding complex host: virus interactions and interference mechanisms will help to develop better interventions and novel anti-viral treatments to prevent infant deaths.

## Methods

### Cell culture and viral propagation

Infant HNO-ALIs were generated as previously described (21, 63). H1 HeLa cell line (ATCC; CAT# CRL-1958) and HEp-2 cell line (ATCC) were grown in minimum essential media (MEM) with 10% fetal bovine serum (FBS) (Corning; CAT# 35-015-CV) and supplemented with 1% L-glutamine (Gibco; CAT# 25030-081) and 1% antibiotic/antimycotic (SIGMA-ALDRICH; CAT# A5955). Human rhinovirus A-16 (RV-A16) strain 11757 purchased from ATCC (CAT# VR-283) was expanded three times in H1 HeLa cells to generate stock viral pools at a consistent and low passage number. Clinical isolate RSV/A/USA/BCM813013/2013(ON) (RSV/A/ON) was expanded three times in HEp-2 cells to establish stock pools and maintain consistent low passage numbers. All cell culture and viral propagation were performed at 36° Celsius/5% CO_2_.

### Sex as a biological variable

Sex was not considered as a biological variable. Infant HNO-ALI lines derived from both female and male donors were utilized (Table S1).

### Coinfection study design

Five HNO-ALI previously derived (21) from healthy infant donors 1. HNO-ALI 9002, 2. HNO-ALI 9003, 3. HNO-ALI 9007, 4. HNO-ALI 90008, 5. HNO-ALI 9009 (Table S1) were used in this study. Two replicate wells per HNO-ALI donor line were infected with 1 of 6 infection conditions: 1) Mock Infected 2) RV-A16 only: multiplicity of infection (MOI) 0.01 3) RSV/A/ON only: MOI 0.01 4) Simultaneous coinfection: RSV/A/ON, MOI 0.01 + RV-A16, MOI 0.01 5. Sequential RV→RSV coinfection: RV-A16, MOI 0.01 then 2 days later RSV/A/ON, MOI 0.01 6. Sequential RSV→RV coinfection RSV/A/ON, MOI 0.01 then 2 days later RV-A16, MOI 0.01. All sequential coinfections occurred two days post initial virus infection (FIG 1A). HNO-ALIs were infected as previously described (21). Samples were collected from the apical compartment as previously described (21, 63) at 8 hours, and 1, 2, 3, 4, 5, 6, 7, 8, 9, and 10 days post infection (dpi). Media from the basolateral compartment was collected and replaced at 2, 4, 6, 8, and 10 dpi.

### Viral quantification

Infectious viral load was determined by quantitative plaque assays. RSV/A/ON was quantified as previously described (21, 63). RV-A16 does not grow in the HEp-2 cell line, so all plaques observed were solely due to the presence of infectious RSV/A/ON. RV-A16 was quantified via 10-fold serial dilutions in MEM/ 2% FBS with 200nM PC786 (MCE CAT#HY-102038), an RSV/A inhibitor, for each sample and plated in duplicate on H1 HeLa monolayers in a 24-well format. After a 1-1.5 hour incubation at 36° Celsius/5% CO_2_ 3.75g methylcellulose +MEM/2%FBS overlay was added to all wells. Plates were further incubated for 3 days at 36° Celsius/5% CO_2_ prior to fixation with 0.01% crystal violet in 10% neutral buffered formalin. The lower limit of detection was 25 PFU/mL for all plaque assays. Any samples with undetectable virus at the 10^-1^ dilution were recorded as half the lower limit of detection (12.5 PFU/mL). RV-A16 and RSV/A/ON positive controls were analyzed alongside every batch for inter-assay consistency and to ensure active inhibition of RSV/A from coinfected samples.

### Histology and immunofluorescence staining and imaging

Histology sections were obtained from HNO-ALIs at 2, 4, and 10 dpi. At each time point one HNO-ALI transwell for each infection condition was fixed using 4% formaldehyde (ThermoFisher; CAT#28908) for 15 minutes followed by successive 30%→50%→70% ethanol dehydration steps at room temperature. HNO-ALI cultures were then cut out, embedded in paraffin, and sectioned. Routine hematoxylin and eosin (H&E) staining was performed. Slides were imaged on ECHO REVOLVE brightfield LED microscope with Olympus 40x/ 0.60NA objective. Representative images displayed are all from the same HNO-ALI donor line (HNO-ALI 9009).

### Lactate dehydrogenase and caspase 3/7 assays

Apical HNO-ALI wash samples at 2, 4, 6, 8, and 10 days post infection (dpi) were assayed for Lactate dehydrogenase (LDH) and caspase 3/7 activity. LDH was quantified using Cytotoxic Detection KitPLUS (Roche Applied Science; CAT#04744934001) according to a previously established protocol (21, 63). Caspase 3/7 was quantified using a previously reported protocol and Caspase-Glo 3/7 kit reagents (Promega; CAT# G8091) as per manufacturer’s established protocol (21, 63).

### Cytokine and chemokine multiplex-immunoassay

Cytokine and chemokines secreted into the apical and basolateral compartment at 2, 4, 6 dpi were measured using 2 separate Milliplex cytokine/chemokine magnetic bead multiplex panels (Millipore). Milliplex Human Cytokine/Chemokine Growth Factor Panel A (Cat# HCYTA-60k-07C) contained human Eotaxin, human interleukin 1 alpha (IL-1α), human interleukin 1 beta (IL-1ß), human interleukin 6 (IL-6), human interleukin 8 (IL-8), human gamma induced protein 10 (IP-10/CXCL10), and human necrosis factor alpha (TNFα). Milliplex Human Interferon Panel (CAT# HIFN-130k-05C) contained human interferon alpha 2 (IFN-α2), human interferon beta (IFN-ß), human interferon lambda 1 (IFN-λ1), human interferon lambda 2 (IFN-λ2), and human interferon lambda 3 (IFN-λ3). All assays were conducted according to manufacturer’s instructions and obtained on a Luminex xPOTENT for MAGPIX v4.2 build 1324 and analyzed using Millliplex Analyst v5.1.0.0 standard build. All analyte concentrations less than the manufacturer’s determined limit of detection for that analyte were considered negative and recorded as half the lower limit of detection.

### Interferon lambda addition assays

Recombinant human IFN-λ obtained from R&D Systems (rhIFN-λ1, CAT#1598-IL-025/CF; rhIFN-λ2, CAT#1587-IL-025/CF; rhIFN-λ3, CAT#5259-IL-025/CF) was diluted in HNO-ALI differentiation media and added to the basolateral compartment of HNO-ALI cultures 24 hours prior to viral infection. Media containing rhIFN-λ was replaced every 48 hours to maintain activity. Apical wash samples were collected at 1, 2, and 3 dpi. RNA was extracted at 3 dpi from all HNO-ALI cultures (Qiagen; CAT# 74106) and reverse transcribed to cDNA (ThermoFisher; CAT#11756500). qPCR for interferon stimulated gene *MX1* (ThermoFisher; CAT#4331182) was performed to confirm IFN activity. Experiments were independently performed in three infant donor lines (HNO-ALI 9002, HNO-ALI 9003, and HNO-ALI 9009). Studies evaluating 10-fold IFN concentrations were independently performed in two infant donor lines (HNO-ALI 9002 and HNO-ALI 9009).

### Interferon lambda receptor 1 blocking assays

To evaluate the role of IFN-λ signaling in restricting RV infection, HNO-ALI cultures were treated with an anti-interferon lambda receptor 1 (IFNLR1) monoclonal antibody (12.5ug/mL; pbl Assay Science, CAT #21885) or isotype control IgG (12.5ug/mL; Cell Signaling, CAT#68860). To model elevated IFN-λ levels present during RSV infection, additional conditions included anti-IFNLR1 or IgG in combination with 100ng/mL rhIFN-λ1 (R&D Systems, CAT#1598-IL-025/CF). Control conditions received either standard HNO-ALI media or 100ng/mL rhIFN-λ1. HNO-ALI differentiation media was used to dilute reagents and added to the basolateral compartment of HNO-ALIs 24 hours prior to infection. Media conditions were refreshed every 72 hours. Each treatment condition was evaluated under four infection conditions: mock infection, RV-A16 only (MOI 0.01), RSV/A/ON only (MOI 0.01), RSV/A/ON→RV-A16 coinfection (each MOI 0.01, 2-day interval as described above). At 5 dpi RNA was extracted (Qiagen; CAT# 74106) and reverse transcribed to cDNA (ThermoFisher CAT#11756500). qPCR for interferon stimulated gene *MX1* (ThermoFisher; CAT#4331182) was performed to verify functionality of anti-IFNLR1 and rhIFN-λ1. Experiments were independently performed in three infant donor lines (HNO-ALI 9002, HNO-ALI 9003, and HNO-ALI 9009).

### Cytokine statistical modeling and analysis

Of the 12 cytokines and chemokines measured, IFN-α2 and IFN-ß were excluded due to zero variance across conditions. Technical replicates were averaged within HNO-ALI line, infection group, and time point prior to analysis. Transformed values were standardized to analyte specific z-scores across all observations and converted to probabilities using the standard normal cumulative distribution function to facilitate cross-analyte comparisons. Analytes were functionally grouped *a priori* into three categories: interferons/antiviral (IFN-λ1, IFN-λ2, and IFN-λ3), chemoattractant (eotaxin and IP-10), and pro-inflammatory (IL-1α, IL-1β, IL-6, IL-8, and TNFα). For each functional group, composite z-scores were calculated as the mean constituent z-score, grouped by the unique combination of HNO-ALI line, infection group, and time point. Composite z-scores were analyzed using generalized linear models with a quasibinomial family and logit link (fractional logit) to assess infection group x dpi interactions while adjusting for surface type (apical vs. basolateral). Post-hoc contrasts comparing infection groups to mock at each time point were estimated as adjusted odds ratio, averaging across surface types. Cluster-robust standard errors accounted for within-HNO-ALI line correlation (Stata v16, glm, yce(cluster)), and results were reported as adjusted odds rations with 95% confidence intervals.

### Statistical analyses and statistical tools

For all area under the curve (AUC) analyses, technical replicates were averaged within HNO-ALI line, experimental group, and time point prior to analysis. Viral replication kinetics were assessed by comparing coinfection AUC with donor-matched single-virus control AUCs, with significance evaluated by using Welch’s t-test across five donor lines. For LDH and caspase 3/7 measurements, infection-condition AUCs were compared with donor-matched mock-control AUCs, and a Welch’s t-test was used to determine statistical significance across infection groups. Association between LDH and caspase were evaluated within each infection group using repeated measures correlation coefficient (*rmcorr*), accounting for clustering of repeated observations within donor lines over time.

AUC analysis was also used to assess the impact of type III IFN treatment on cumulative viral load. Within each donor line, IFN-treated AUCs were compared with corresponding untreated virus controls, and significance determined using a Welch’s t-test across three donor lines. When testing 10-fold concentrations of type III IFNs, a two-way analysis of variance (ANOVA) was performed to evaluate significant impacts on RV replication. All graphs were made using Graphpad Prism v11.02.

Unless otherwise stated, all statistical analyses were conducted in R v 3.19 (R Core Team, 2024) using the following packages: *lme4* (*64*), *lmerTest* (65), *nlme* (66)*, rmcorr* (*67*)*, ggplot* (*68*)*, ggpubr* (*69*)*, RColorBrewer* (*70*)*, sandwich* (71), and *tidyr* (*72*). Statistical significance was assessed at the alpha = 0.05 level for all tests.

### Study approval

Nasal washes and swabs to generate nasal organoids were previously collected from infants after obtaining signed informed consent from the adult or legal guardian under an approved protocol by the Institutional Review Board at Baylor College of Medicine (21).

### Data availability

A supporting data values file with all reported data values will be available as part of the Supplemental Material.

## Author Contributions

EMS conceptualized the project, performed experiments, analyzed data, obtained funding, and wrote the original manuscript draft. GMA, AM, SW, LA, TM, NO, HL, GK, DN helped perform experiments and edited the manuscript. DH and LFS performed statistical modeling and edited the manuscript. SEB supervised established organoid cultures, obtained funding, and edited the manuscript. PAP and VA conceptualized the project, analyzed data, obtained funding, and edited the manuscript. All authors reviewed and approved the final manuscript.

## Acknowledgments

We would like to thank Alejandra Rivera Tostado, Xi-Lei Zeng, Aaya I. Boussattach, and Mia Borrego in the Baylor College of Medicine 3D Organoid Core for their collaboration in maintenance of established HNO-ALI donor lines. Experimental schematic images were created using BioRender.com.

## Supplemental Figure Legends

**Supplemental Figure S1. Individual cytokine or chemokine release in different coinfection sequences across *time.*** Forest plots of the adjusted odds ratio with 95% confidence interval for individual cytokines tested. Comparisons between different infection groups at 2, 4, and 6 dpi for **(A)** IFN-λ1 **(B)** IFN-λ2 **(C)** IFN-λ3 **(D)** Eotaxin **(E)** IP-10 **(F)** TNFα **(G)** IL-1α **(H)** IL-1ß **(I)** IL-6 **(J)** IL-8. Data analyzed from n=2 technical replicates from five infant donor lines. Significance indicated by non-overlapping confidence intervals. See also Supp Data.

**Supplemental Figure S2. Type III interferon expression heterogeneity between different infant donor HNO-*ALI lines.*** Comparison of IFN-λ1, -λ2, and -λ3 expression at 2, 4, and 6 dpi with either RV or RSV infection between HNO-ALI donor lines **(A)** 9002 **(B)** 9003 **(C)** 9007 **(D)** 9008 **(E)** 9009. Data displayed as individual n=2 technical replicates.

**Supplemental Figure S3. Type III interferons inhibit rhinovirus replication but not RSV replication (A)** RV-A16 replication over time in the presence of type III interferons (IFN) **(B)** RSV/A/ON replication over time in the presence of type III IFNs. All data displayed as average±SD of n=2 technical replicates from 3 different infant donor lines. Dotted line indicates limit of detection (LOD). *Not Significant (N.S.),* \**p<0.05, **p<0.01, ***p<0.*001 by Two-way ANOVA. **(C-E)** Validation of functional type III interferons by qPCR *MX1* expression at 5dpi for individual infant HNO-ALI donor lines. Individual technical replicates shown.

**Supplemental Figure S4. Different type III interferons have similar inhibition of rhinovirus replication at *multiple concentrations.*** RV-A16 replication over time in the presence of either rhIFN-λ1, rhIFN-λ2, or rhIFN-λ3 at **(A)** 100ng/mL **(B)** 10ng/mL or **(C**) 1ng/mL. All data displayed as average±SD of n=2 technical replicates from 2 different infant donor lines. Dotted line indicates limit of detection (LOD). **(D-E)** Validation of functional type III interferons by qPCR *MX1* expression at 5 dpi for individual infant HNO-ALI donor lines. Individual technical replicates shown.

**Supplemental Figure S5. Monoclonal antibody against IFNLR1 blocks MX1 expression in infant HNO-ALIs. (A, C, E)** Validation of functional mAb anti-IFNLR1 as measured by induction of interferon stimulated gene *MX1* in all experimental groups 5 days post infection **(B, D, F)** Validation of functional mAb anti-IFNLR1 and IFN-λ1 as measured by *MX1* induction 5 days post infection. Three different donor lines were utilized in these experiments **(A, B)** HNO-ALI 9002 **(C, D)** HNO-ALI 9003 **(E, F)** HNO-ALI 9009.

## Supplemental Table

**Supplemental Table S1.** Donor demographics for infant-derived HNO-ALIs.

| BCM Internal Line<br>Identification<br>Number | Age at collection | Sex | Race/Ethnicity | Health Status |
| --- | --- | --- | --- | --- |
| HNO-ALI 9002 | <24 months | Female | Asian and White, Non-Hispanic | Healthy |
| HNO-ALI 9003 | <12 months | Female | White, Non-Hispanic | Healthy |
| HNO-ALI 9007 | <12 months | Female | Asian and White, Non-Hispanic | Healthy |
| HNO-ALI 9008 | <24 months | Male | White, Non-Hispanic | Healthy |
| HNO-ALI 9009 | <12 months | Male | White, Non-Hispanic | Healthy |

